# *In vivo* structure probing of RNA in *Archaea*: Novel insights into the ribosome structure of *Methanosarcina acetivorans*

**DOI:** 10.1101/2023.04.14.536875

**Authors:** Allison M. Williams, Elizabeth A. Jolley, Michel Geovanni Santiago-Martínez, Cheong Xin Chan, Robin R. Gutell, James G. Ferry, Philip C. Bevilacqua

**Affiliations:** Department of Biochemistry and Molecular Biology, Pennsylvania State University, University Park, PA, 16802, United States; Center for RNA Molecular Biology, Pennsylvania State University, University Park, PA, 16802, United States; Department of Chemistry, Pennsylvania State University, University Park, PA, 16802, United States; Australian Centre for Ecogenomics, School of Chemistry and Molecular Biosciences, The University of Queensland, Brisbane QLD 4072, Australia; Department of Integrative Biology, The University of Texas at Austin, Austin, TX, 78712, United States; Division of Rheumatology, Inflammation, and Immunity, Department of Medicine Brigham and Women’s Hospital, Harvard Medical School, Boston, MA 02115, United States; Department of Molecular and Cell Biology, The University of Connecticut, Storrs, CT, 06269, United States

## Abstract

Structure probing combined with next-generation sequencing (NGS) has provided novel insights into RNA structure-function relationships. To date such studies have focused largely on bacteria and eukaryotes, with little attention given to the third domain of life, archaea. Furthermore, functional RNAs have not been extensively studied in archaea, leaving open questions about RNA structure and function within this domain of life. With archaeal species being diverse and having many similarities to both bacteria and eukaryotes, the archaea domain has the potential to be an evolutionary bridge. In this study, we introduce a method for probing RNA structure *in vivo* in the archaea domain of life. We investigated the structure of ribosomal RNA (rRNA) from *Methanosarcina acetivorans*, a well-studied anaerobic archaeal species, grown with either methanol or acetate. After probing the RNA *in vivo* with dimethyl sulfate (DMS), Structure-seq2 libraries were generated, sequenced, and analyzed. We mapped the reactivity of DMS onto the secondary structure of the ribosome, which we determined independently with comparative analysis, and confirmed the accuracy of DMS probing in *M. acetivorans*. Accessibility of the rRNA to DMS in the two carbon sources was found to be quite similar, although some differences were found. Overall, this study establishes the Structure-seq2 pipeline in the archaea domain of life and informs about ribosomal structure within *M. acetivorans*.

## Introduction

Appreciation of the role of RNA structure in biology has greatly expanded over the last decade, in large part because of the development of next generation sequencing (NGS) (Kwok et al. 2015; Qian et al. 2019; Zhao et al. 2019). RNA has emerged as an essential biopolymer that allows organisms to respond to their surroundings (Kavita and Breaker 2023). In combination with biophysical insight into RNA, NGS has advanced our understanding of the role of RNA structure in cellular function on a massive scale. Over the last decade, NGS tools have been developed that permit investigation of RNA structure *in vivo* utilizing diverse chemical probes (Ritchey et al. 2017; Feng et al. 2018; Busan et al. 2019; Mitchell et al. 2019a, 2019b; Wang et al. 2019).

To date, *in vivo* RNA structure has been probed transcriptome-wide in multiple organisms and viruses, including but not limited to yeast (Rouskin et al. 2014), mammalian cells (Waldron et al. 2019; Piao et al. 2022), gram negative bacteria (Incarnato et al. 2017), gram positive bacteria (Ritchey et al. 2020), SARS-CoV-2 (Manfredonia et al. 2020), a dicot plant (Ding et al. 2014; Tack et al. 2020), and a monocot plant (Su et al. 2018). These investigations have revealed the nature of RNA structure under diverse growth conditions including unstressed (Ding et al. 2014), amino acid-stressed (Ritchey et al. 2020), heat-shocked (Su et al. 2018) heat-varied (Jolley et al.), and salt-stressed growth conditions (Tack et al. 2020). In addition to the extensive research on eukaryotic and bacterial systems, the archaea domain of life has been explored with respect to RNA half-life, abundance, and *in vivo* interactions (Peterson et al. 2016; Knüppel et al. 2020; Gehlert et al. 2022). Nonetheless, methods for probing RNA structure have not yet been widely adapted or applied to archaea. Such approaches stand to elucidate the function of RNA structure in archaea and to expand the role of RNA structure across all three domains of life.

*Methanosarcina acetivorans* is a model methane-producing archaeon and an optimal candidate for *in vivo* structure probing (Ferry 2020). Study of *M. acetivorans* facilitates mechanistic understanding of the conversion of acetate to methane, which accounts for most of the 400-500 Tg of biogenic methane produced annually effecting global warming and climate change (Conrad 2009). Indeed, Earth’s greatest mass extinction in the end-Permian carbon cycle is credited to ancestors of *M. acetivorans* that evolved the pathway for converting acetate to methane producing a methanogenic burst and a rapid increase in global warming (Rothman et al. 2014). Moreover, *M. acetivorans* contains a larger and more diverse genome than many other archaea; for instance, its genome size is 5.75 Mbp, while the closely related *M. mazei* has a genome size of just over 4 Mbp (Maeder et al. 2006). Additionally, *M. acetivorans* has distinct metabolic pathways for carbon processing, allowing it to grow with methanol or acetate (Galagan et al. 2002). These two carbon and energy sources lead to different growth rates, as well as activation of different genes and mRNA half-lives (Li et al. 2007; Peterson et al. 2016). Moreover, these two types of growth substrates could present unique intracellular conditions for RNA folding. Some genes are differentially expressed between these two growth conditions, while housekeeping genes, including those for ribosome biogenesis, are expressed constitutively, albeit in different quantities (Li et al. 2007; Peterson et al. 2016). Ribosome structures from different archaea have been studied by X-ray crystallography and cryo-EM, including *Haloarcula marismortui* (Ban et al. 2000; Gabdulkhakov et al. 2013), *Methanothermobacter thermautotrophicus* (Greber et al. 2012), *Pyrococcus furiosus* (Armache et al. 2013), *Pyrococcus abyssi* (Coureux et al. 2020), and *Thermococcus celer* (Nürenberg-Goloub et al. 2020). However, archaeal ribosome structures have not been probed chemically *in vivo*, and potential differences in ribosome structure with respect to carbon and energy sources are unknown.

In this study, we present an experimental approach to probe RNA structure *in vivo* in a model archaeon that results in high quality RNA samples ready for NGS library preparations. Our approach uses Structure-seq2 libraries (Ritchey et al. 2017; Tack et al. 2018) for *M. acetivorans* grown with either methanol or acetate and probed *in vivo* with dimethyl sulfate (DMS), which is A- and C-specific (Mitchell et al. 2019a). To validate the approach with DMS probing, we focused herein on data pertaining to rRNA. We used comparative analysis to establish the secondary structures of the 5S, 16S and 23S rRNA in *M. acetivorans* and then mapped our *in vivo* probing data onto these structures under both growth conditions. Chemical mapping data were largely consistent with these comparative structures, validating our *in vivo* probing. Chemically accessible regions in the rRNA were very similar between the two growth conditions, although some differences were found. Overall, this study sets the stage for the use of Structure-seq2 transcriptome-wide in archaea, with the potential for discovery of novel regulatory RNAs.

## Materials and Methods

### DNA oligonucleotides

Sequences for all DNA oligonucleotides used in this study are provided in Supplemental Table 1. Oligonucleotides were ordered from Integrated DNA Technologies (IDT) and purified as noted.

### Growth of Methanosarcina acetivorans

*M. acetivorans* C2A (DSM 2834, ATCC 35395) was cultured under anoxic conditions in high salt (HS) media (pH 6.8) supplied with 100 mM acetate or 100 mM methanol as energy and carbon sources, as previously reported. A detailed protocol is described in (Santiago-Martínez and Ferry 2023). Briefly, Milli Q water was placed into an anaerobic chamber (COY laboratory products, Grass Lake, Michigan, USA) with 80% N_2_, 15% CO_2_ and 5% H_2_. Then, the following salts were added: 23.4 g/L NaCl, 3.8 g/L NaHCO_3_, 1.0 g/L KCl, 11 g/L MgCl_2_, 0.2 g/L CaCl_2_, 1% (v/v) vitamin solution, 1% (v/v) trace mineral solution, and 0.001% (w/v) resazurin as redox indicator. The medium was bubbled with the mix of gases described above overnight. Next, 1.0 g/L NH_4_Cl (nitrogen source) and 0.5 g/L cysteine-HCl (to ensure complete chemical reduction) were added. Then, 100 mL of medium were poured into 120 mL serum bottles, sealed with butyl rubber stoppers, and secured with aluminum crimp caps. The medium was autoclaved at 121 °C for 30 min. Before cell inoculation, 10 mL/L of 2.5 % Na_2_S•9 H_2_O and 5.0 mL/L of 1 M KH_2_PO_4_ were added from anaerobic sterile stock solutions, as well as 100 mM methanol or acetate. Cultures were started by adding cell inoculum (which proceeded from cell cultures grown with methanol or acetate) to fresh HS medium for further incubation at 37°C without shaking. Cells were grown independently in biological triplicate, and growth was monitored by changes in Optical Density (OD) at 600 nm. Cells were harvested in mid-exponential phase (OD_600_ = 0.4 for methanol and OD_600_ = 0.2 for acetate) (Lira-Silva et al. 2012) and used for RNA extractions as diagrammed in Supplemental Fig. S1.

### In vivo DMS structure probing

Treatment of cells with DMS was adapted from our previous protocol (Ritchey et al. 2020). All DMS treatments were conducted in a chemical fume hood for safety, under aerobic conditions. Either 6 or 12 mL of methanol-or acetate-grown cells, respectively, were centrifuged at 5000 rpm (4300 xg) for 20 min at 4 °C (Supplemental Fig. S1). The volume of acetate-grown cells was larger to have equivalent biomass because at exponential phase, the OD_600_ for acetate is half that in methanol. Pellets were resuspended in 1 mL of HS media and incubated with and without 150 mM DMS for 3 min with occasional mixing. Tubes were then tightly capped and centrifuged at 7000 rpm (8400 xg) for 5 min at 4 °C. Supernatant was decanted into excess dithiothreitol (DTT) quench solution. Notably, it was important to not add DTT directly to the cells (see Results). Samples +/– DMS were resuspended in 1 mL of Trizol, and the RNA was extracted according to the Direct-zol RNA MiniPrep kit (Zymo Research). This treatment provided rRNA that is within single hit kinetics range (Supplemental Fig. S2) (Tijerina et al. 2007; Kertesz et al. 2010; Wan et al. 2011; Singulani et al. 2017).

### Library preparation using Structure-seq2

After total RNA extraction, RNA quality was assessed as prokaryote total RNA on an Agilent 4150 TapeStation (Penn State Genomics Facility) and found to have favorable RIN scores (≥ 8.5). Subsequently, libraries were prepared as described previously (Ritchey et al. 2017). Twelve libraries were generated: six with methanol-grown cells (three biological replicates, each +/– DMS) and six with acetate-grown cells (three biological replicates, each +/– DMS). Briefly, 500 ng of total RNA was reverse transcribed using SuperScript III, incorporating a minor fraction of biotinylated dCTP and dUTP along with the standard dNTPs. Samples were passed through an RNA Clean & Concentrator kit (Zymo Research) to remove proteins and free nucleotides, and then mixed with 25 µL of streptavidin magnetic beads to collect the biotinylated cDNA, which was purified with a second RNA Clean & Concentrator. Hairpin adaptors were ligated onto the 3′-end of reverse transcription products (Kwok et al. 2013), and the cDNA was purified through a third RNA Clean & Concentrator. A second streptavidin magnetic bead pulldown and a fourth RNA Clean & Concentrator step were completed followed by a PCR cycle test to identify the lowest cycle number of cycles (= 25) that created a visible product on an agarose gel. The amplified libraries were fractionated on an 8.3 M urea, 10 % polyacrylamide gel and size-selected between 200 and 600 nt. Samples were recovered by an overnight crush and soak and ethanol precipitation. Samples were submitted to the Penn State Genomics Core Facility to confirm uniformity of the smear on the Agilent TapeStation 4150. Sequencing was performed at the same facility using 150 nt single-end sequencing on a NextSeq 2000 Mid Output platform.

### DMS reactivity analyses and quality assessment

To assess and analyze the library reads, we followed the StructureFold2 pipeline (Tack et al. 2018). Briefly, samples were trimmed with Cutadapt v1.16 and mapped to the transcriptome NC_003552.1 that contained RNA transcripts with no untranslated regions, allowing multimapping (Supplemental Table 2). These samples were filtered with the default parameters in StructureFold2 using the sam_filter module, which removed reads with first-base mismatches, reads with greater than three mismatches, and reads that mapped to the reverse strand. Downstream analysis was performed on only one set of rRNA genes (locus tags MA_RS04665-16S rRNA, MA_RS04670-23S rRNA, and MA_RS04675-5S rRNA). This was done for simplicity as each copy of a specific rRNA gene differs from the other copies by no more than 2% nucleotide identity. Mapped reads were converted to reverse transcriptase stop count (RTSC) files, which were correlated using the raw RT stops to ensure confidence among the three replicates. All replicates were found to be well correlated and so were pooled within each of the four conditions (acetate/methanol X +/– DMS). The DMS RTSCs were converted to DMS reactivities utilizing the rtsc_to_react module in StructureFold2 and normalized for the number of RT stops per nucleotide and the length of the transcript (Tang et al. 2015). DMS reactivity was manually superimposed onto the appropriate secondary structure. We used 0.2 as the minimum cutoff for mapping DMS reactivity on the secondary structure for each condition as per Yamagami et al. (2022) and 0.5 as the minimum DMS reactivity for differences.

### Determination of the M. acetivorans rRNA secondary structure with comparative analysis

The *M. acetivorans* 5S, 16S, and 23S rRNA secondary structures were determined with comparative analysis. This method, first applied to the tRNA cloverleaf secondary structure, is based on the simple realization that all the sequences within any given RNA family will form the same RNA secondary and three-dimensional structure. Due to the structural equivalence of the six canonical Watson-Crick and wobble base pair types, different RNA sequences can interchange these base pair types within the conserved helices to maintain the same RNA structure. While different computational methods can identify a common structure for all the sequences in each RNA family, the preferred method here is based on the search for positions (columns) within a sequence alignment that have the same pattern of variation, or covariation. Base pairs are inferred as being between those positions that covary. Several covariation algorithms were developed and used to determine the comparative structures for tRNA and for 16S and 23S rRNAs (Gutell et al. 1992; Cannone et al. 2002). After more than twenty years of developing and refining the16S and 23S rRNA covariation-based secondary structures, their accuracy was accessed when several high-resolution crystal structures of the ribosome were determined (Ramakrishnan 2002). Astoundingly, ∼98% of the base pairs with covariation were present in the crystal structures (Gutell et al. 2002).

The determination of the comparative covariation-based rRNA structures has evolved in stages. Earlier stages were establishing higher quality structure-based sequence alignments, the most effective covariation algorithms, and the detailed procedures for sequence alignment and the interpretation of the results from the covariation analysis. Current analysis utilizes the well curated structure-based alignments with a large set of sequences that contain significant diversity in sequences and structures with a broad phylogenetic sampling, and mature covariation-based secondary structure models (Gutell et al. 1985, 1994; Gutell 2014).

The *M. acetivorans* rRNA sequences were manually aligned into the set of previously aligned rRNA sequences using the Jalview alignment editor (https://www.jalview.org/). This structure-based sequence alignment procedure assured the most accurate alignment while simultaneously determining the *M. acetivorans* secondary structure during the alignment process. Secondary structure diagrams were templated off previously generated archaeal 5S, 16S, and 23S rRNA secondary structure diagrams available at the CRW Site (https://crw2-comparative-rna-web.org/database/) (Cannone et al. 2002). Specifically, *M. acetivorans* 5S rRNA was templated from *Methanobacterium formicicum*, 16S rRNA was templated from *Methanococcus vannielli*, and the 23S rRNA was templated from *Haloarcula marismortui*. After templating, the *M. acetivorans* secondary structure diagrams were refined with the XRNA secondary structure editor (http://rna.ucsc.edu/rnacenter/xrna/xrna.html). The layout of the helices in the *M. acetivorans* 5S, 16S and 23S rRNA secondary structure diagrams are similar to those originally established by Noller and Woese (Fox and Woese 1975; Noller and Woese 1981; Noller et al. 1981) and available at the CRW Site (vide supra). Helices are numbered according to Brimacombe (1995).

## Results

### In vivo RNA structure probing in M. acetivorans

Due to the unique physiology of *M. acetivorans*, special consideration was taken while developing a method for probing RNA structure *in vivo*. In particular, this archaeon has a membrane layer called the “S-layer”, which is comprised of proteins, stabilized by magnesium ions, and sensitive to reducing solutions (Arbing et al. 2012). As a result, several steps were specialized for the purpose of extracting high quality, DMS-treated RNA from *M. acetivorans*. Notably, owing to the unique composition of this S-layer, it was important to not add DTT directly to the cells, as was done previously for bacteria and eukaryotes (Ritchey et al. 2020).

Methods development for *in vivo* probing have often examined rRNAs, as they are abundant and can be accurately modeled by comparative approaches (Ding et al. 2014; Mitchell et al. 2017, 2019b), and therefore the Structure-seq2 libraries were not depleted of rRNA. We chose DMS for developing this method in *Archaea* because it is the most widely used chemical probe and has been used to study a wide range of systems and stresses (Bevilacqua and Assmann 2018; Mitchell et al. 2019a) The libraries were of high quality with 50 to 86% of the effective reads mapping to the transcriptome (Supplemental Table 2). To assess the nucleobase-specificity of DMS reactivity, we calculated percentage of RT stops at A and C residues. For the methanol and acetate –DMS samples, percent stops were relatively evenly distributed across the four nucleotides in all three biological replicates as expected (Supplemental Table 3). In the +DMS samples, percent stops for A increased from ∼23% to ∼60% for both methanol and acetate samples. The RT stop percentages for Cs were second highest for acetate samples, although not for methanol samples. Significantly lower methylation of C by DMS has been seen for previous Structure-seq experiments (Ding et al. 2014; Su et al. 2018; Ritchey et al. 2020). These observations support DMS penetrating *M. acetivorans* cells and methylating RNA in a sequence-specific manner.

To assess reproducibility, the pairwise combinations of the three biological replicates were compared for each treatment under each growth condition (Supplemental Figs. 3 and 4). The replicates were well correlated, with r-values ≥ 0.85, and therefore pooled. The DMS reactivity was calculated on pooled triplicate samples. In sum, we demonstrated the ability to probe *M. acetivorans* RNA with DMS *in vivo* in a base-specific manner and to create high quality libraries that were mappable and well-correlated across replicates. Next, we wanted to test if DMS reactivity was consistent with RNA structure. To do so, we first determined the comparative structure of the rRNA.

### The M. acetivorans 5S, 16S, and 23S rRNA comparative secondary structures

We determined comparative secondary structures for *M. acetivorans* 5S, 16S and 23S rRNAs structures. These are provided in Fig. 1 and Supplemental Fig. S5 through S13, which show base pairing and labeling in standardized formats according to the *E. coli* ribosome. Comparison of the *M. acetivorans* comparative secondary structure with the archaeal rRNA conservation secondary structure diagrams^1^ revealed, as expected, that the *M. acetivorans* rRNAs had sequence and structural features characteristic of the archaea. Indeed, the *M. acetivorans* rRNA sequences and secondary structures superimposed, without any exceptions or anomalies, onto 5S, 16S and 23S rRNAs secondary structures that were previously determined to be ∼98% accurate. Thus, we can extrapolate with confidence that the base pairs in the *M. acetivorans* 5S, 16S and 23S rRNAs covariation-based secondary structure models presented herein are correct. One can also be confident that the conformation for nearly all of the depicted base pairs is the standard Watson-Crick three-dimensional conformation (Lee and Gutell 2004; Stombaugh et al. 2009). However, while the covariation analysis accurately identifies base pairs (and these base pairs nearly always form the standard Watson-Crick conformation – cWW), it should be noted that many of the nucleotides that form the hairpin, internal, and multi-stem loops in these covariation-based structure models are not unpaired in the crystal and cryo-EM three-dimensional structures (Lee and Gutell 2004; Stombaugh et al. 2009) where they form myriad non-cWW base pair conformations.

**Figure 1:**
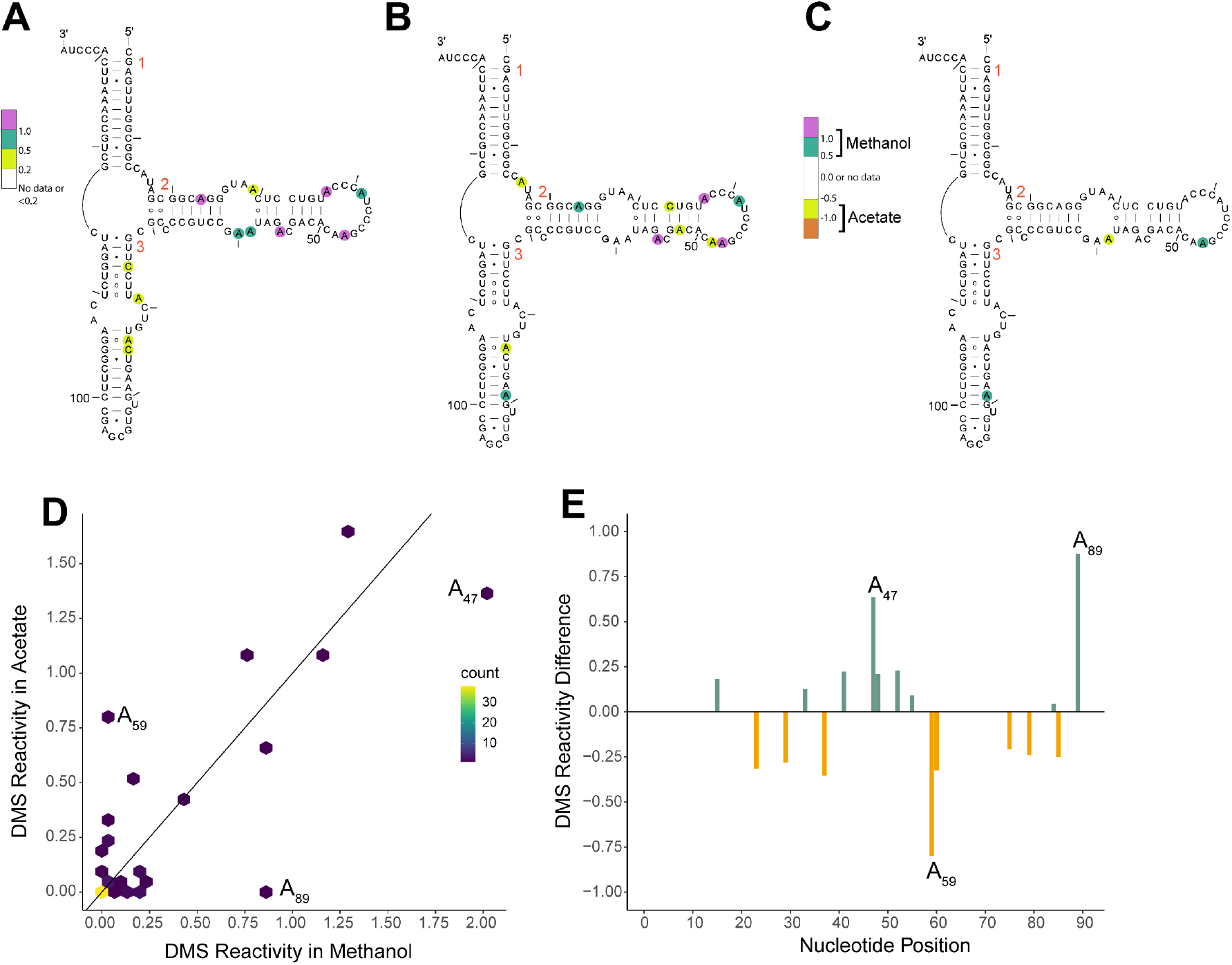
Comparison of 5S rRNA DMS reactivity in methanol and acetate. DMS reactivity in **(A)** acetate, **(B)** methanol, and **(C)** methanol – acetate. Colored A and C nucleotides have significant DMS reactivity (see legends). Helices are numbered according to standardized format and nucleotides have a tick every 10 nucleotides. **(D)** Correlation plot of acetate and methanol DMS reactivity. Line indicates y=x. **(E)** DMS reactivity difference plot. Each bar was calculated as DMS reactivity in methanol – acetate. Green bars indicate higher DMS reactivity in methanol, whereas orange bars indicate higher DMS reactivity in acetate. Select bars are annotated for reference.

### DMS reactivity aligns with comparative ribosomal structure

With comparative structures established for the three *M. acetivorans* rRNAs, we assessed whether DMS reactivity is consistent with them according to the steps described in the Materials and Methods. For the ease of comparison to other organisms, we have incorporated the equivalent nucleotide position in *E. coli* in parentheses. We first consider 5S rRNA in acetate-grown *M. acetivorans* cells where DMS reactivity mapped to bulges and to internal and hairpin loops (Fig. 1A). For instance, A_37_ (C_35_), A_47_ (A_45_), and A_55_ (A_53_) were highly reactive and found in such positions. For the 5S rRNA in methanol-grown samples, the DMS reactivity mapped to the same three A residues and to very similar positions overall (Fig. 1B). This is further evident when the differences in reactivity between methanol and acetate conditions are mapped onto the 5S rRNA secondary structure (Fig. 1C). Only three nucleotides show differences in reactivity of 0.5 or greater, with two of them (A_47_ (A_45_) and A_89_ (G_83_) greater in methanol and one (A_59_ (A_57_)) greater in acetate. Both A_47_ (A_45_) and A_59_ (A_57_) are in loop regions, while A_89_ (G_83_) is in a helical region. To visualize the correlation between conditions, we plotted the DMS reactivity of acetate-vs. methanol-grown samples (Fig. 1D). As expected, the majority of the reactivities were relatively similar between the two media. There were a few nucleotides that had high reactivities, but these were similar in both media as indicated by the presence of points near the end of the diagonal line. Similarity in reactivity is also evidenced by the DMS reactivity difference bar plots, which showed no data with |DMS reactivity differences| > 1.0. The three A nucleotides mentioned above with reactivities between 0.5 and 1.0 are annotated (Fig. 1E). It is notable that two of these positions, A_59_ (A_57_) and A_89_ (G_83_) showed strong reactivity in one condition but almost none in the other (Fig. 1D). The underlying reasons for differential reactivity in one growth-medium over the other could be due to preferential protein binding in one media binding but is presently unclear. Changes in the relative abundance of several yeast ribosomal proteins has been observed by both mass spectrometry and cryoEM upon shifting the growth media from glucose to glycerol, supporting this general possibility (Samir et al. 2018; Sun et al. 2021).

We repeated this analysis for the 16S rRNA. DMS reactivity for both acetate- and methanol-grown samples again mapped largely to bulges and to internal and hairpin loops, particularly for highly reactive nucleotides (Supplemental Fig. S5 and S6). One exception occurs at helix 44 (Fig. 2A), where many of the nucleotides exhibited DMS reactivity despite base pairing. In acetate, DMS reactivity is found across 33% of the helix, while in methanol DMS reactivity occurs across 20% of the helix. Another region of interest is helix 27 (Fig. 2B). In both growth conditions, the internal loop of the helix shows reactivity for all A and C nucleotides, indicating this is a region of high chemical accessibility. To better understand potential differences in reactivity between the two media, we plotted the DMS reactivity of acetate-versus methanol-grown samples (Fig. 2C), as well as differences in reactivity (Fig. 2D and Supplemental Fig. S7 mapped to structure). The majority of the reactivities in the two growth media were relatively similar to each other, as in 5S rRNA. There were a few nucleotides that have high reactivities above 2.0, but these were similar in both media as indicated by the presence of points near the end of the diagonal line (Fig. 2C). There were four nucleotides that had differential DMS reactivity near +1 or –1 in 16S rRNA (Fig. 2D and Supplemental Fig. S7). Two of these, C_784_ (C_841_) and C_1080_ (C_1137_), were more reactive in methanol, while two others, C_474_ (U_534_) and C_1358_ (C_1411_), were more reactive in acetate. Except for C_1358_ (C_1411_), these nucleotides are in single stranded regions, consistent with accessibility to chemical probing. Residue C_1358_ (C_1411_) is part of helix 44, the highly reactive region described earlier. As evidenced by the lack of colored nucleotides flanking these four residues (Supplemental Fig. S7), both media showed similar DMS reactivity in these four regions indicating a lack of a global conformational change. It is notable that three of these positions showed strong reactivity in one condition but almost none in the other (Fig. 2C). Again, the basis for these media-specific differences in rRNA DMS reactivity is unclear but could be due to preferential protein binding in one media.

**Figure 2:**
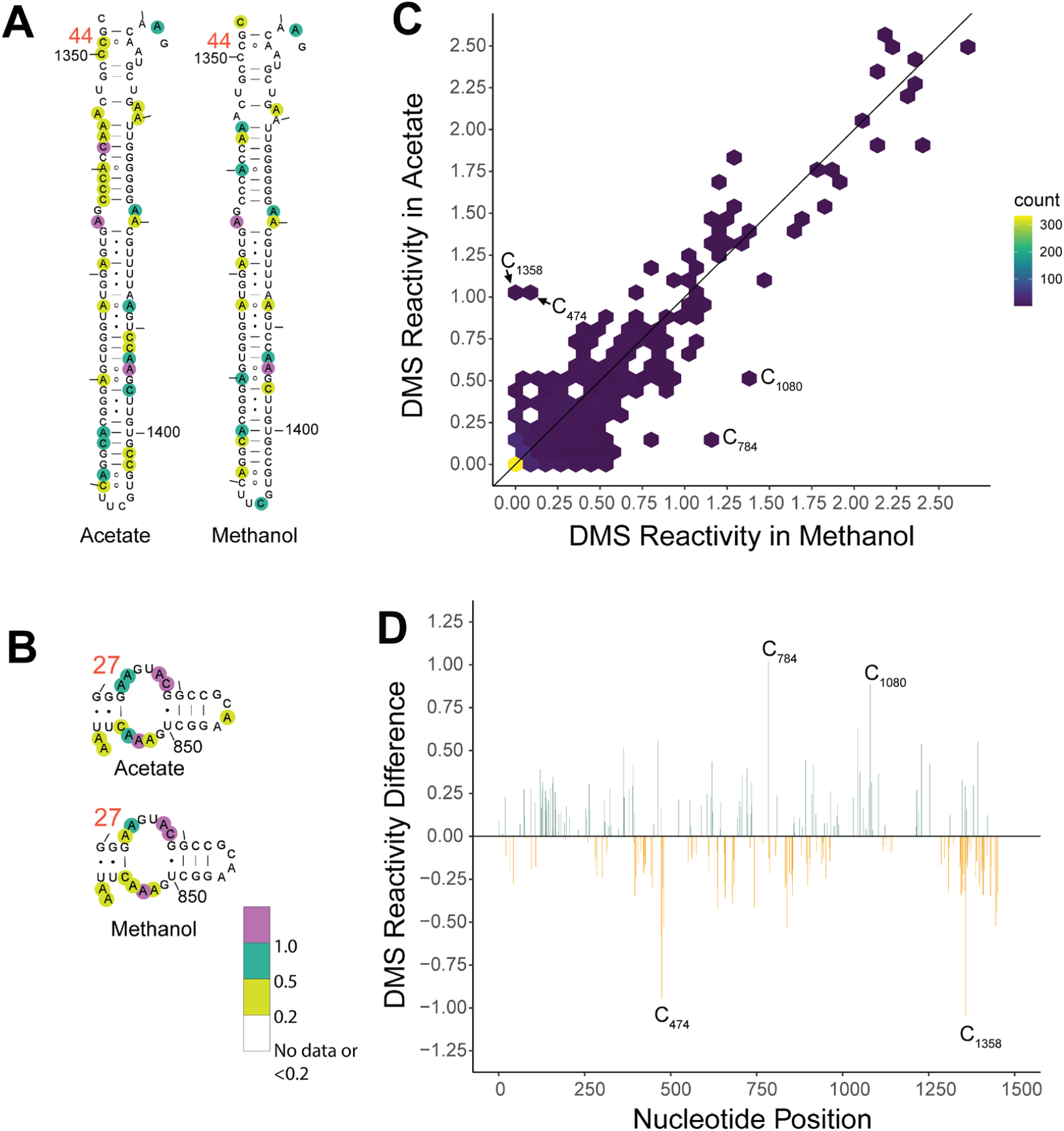
Comparison of 16S rRNA DMS reactivity in methanol and acetate. DMS reactivity comparison in acetate and methanol for **(A)** Helix 44 and **(B)** Helix 27. Both panels are oriented as in the full 16S rRNA diagram found in the Supplemental Information. Colored A and C nucleotides have significant DMS reactivity (see legends). Helices are numbered according to standardized format and nucleotides have a tick every 10 nucleotides. **(C)** Correlation plot of acetate and methanol DMS reactivity. Line indicates y=x. **(D)** DMS reactivity difference plot. Each bar was calculated by DMS reactivity in methanol – acetate. Green bars indicate higher DMS reactivity in methanol, whereas orange bars indicate higher DMS reactivity in acetate. Select bars are annotated for reference.

Finally, we compared DMS reactivity within the 23S rRNA. As observed for 5S and 16S rRNA, DMS reactivity for acetate samples and methanol samples again largely mapped to bulges and to internal and hairpin loops (Supplemental Figs. S8, S9, S11, S12). For instance, stems 43, 43.1 and 44 (Fig. 3A) contain several highly reactive nucleotides, above 0.5, almost all of which are in loops. To better understand potential differences in reactivity between the two media, we plotted the DMS reactivity of acetate-versus methanol-grown samples (Fig. 3B), as well as differences in reactivity (Fig. 3C and Supplemental Figs. S10 and S13 mapped to the structure). Again, most reactivities in the two growth media were similar to one another including some highly reactive positions indicated by the presence of points near the end of the diagonal line (Fig. 3B). There were two nucleotides with DMS reactivity differences near +1 or –1 in 23S rRNA: in methanol, A_1578_ (C_1488_), which is involved in an A•G base pair in helix 58 and in acetate, A_873_ (A_783_), which is single-stranded in helix 35a. As evidenced by the lack of colored nucleotides flanking these two residues (Supplemental Fig. S10), both media showed similar DMS reactivity overall in these two regions of the 23S rRNA, indicating similar structures overall. Again, the basis for these media-specific differences in rRNA DMS reactivity is unclear but could be due to preferential protein binding in one media.

**Figure 3:**
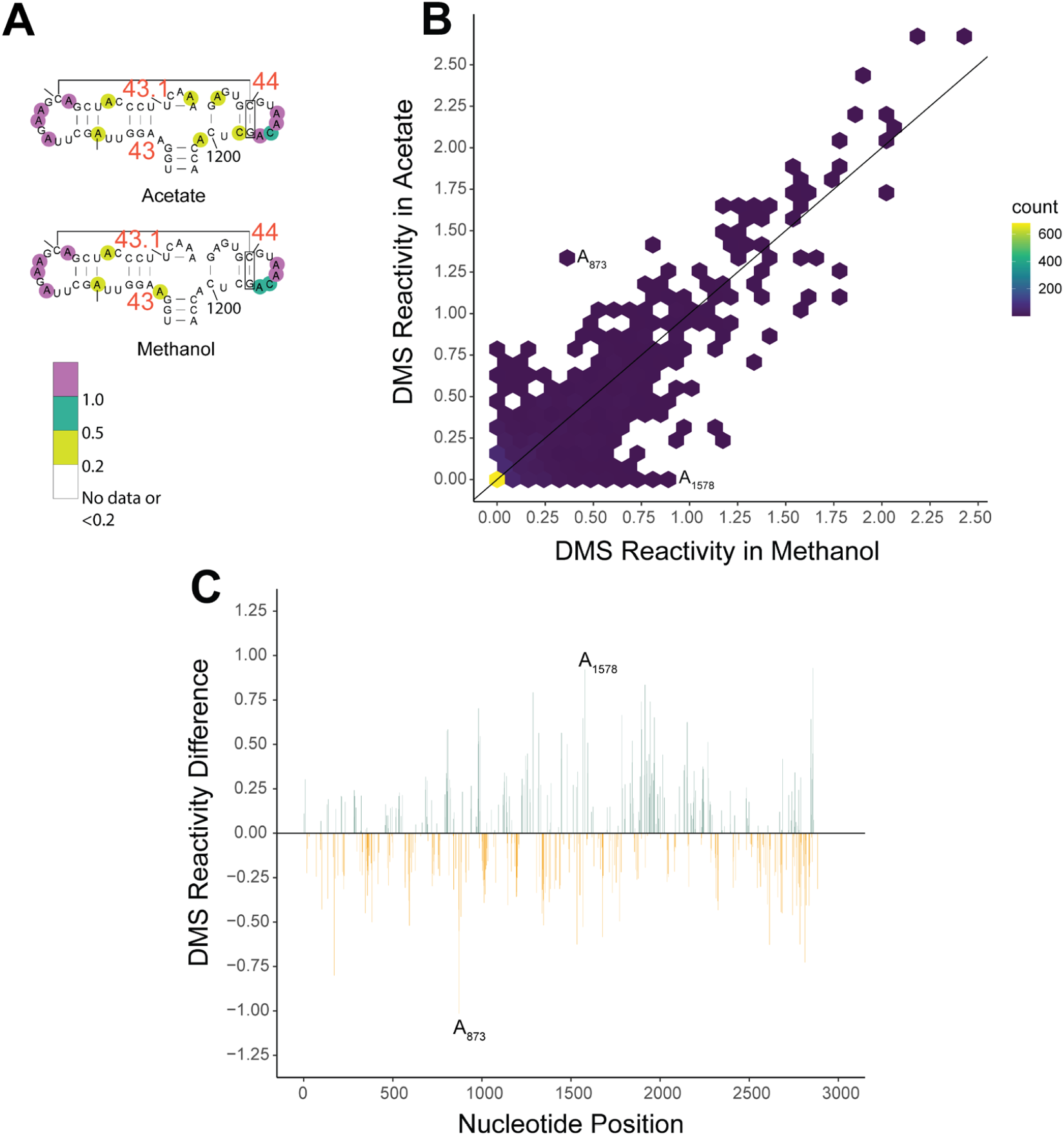
Comparison of 23S rRNA DMS reactivity in methanol and acetate. DMS reactivity comparison between acetate and methanol for **(A)** Helices 43, 43.1, and 44. Panel is oriented as in the full 23S rRNA diagram found in the Supplemental Information. Colored A and C nucleotides have significant DMS reactivity (see legends). Helices are numbered according to standardized format and nucleotides have a tick every 10 nucleotides. **(B)** Correlation plot of acetate and methanol DMS reactivity. Line indicates y=x. **(C)** DMS reactivity difference plot. Each bar was calculated by DMS reactivity in methanol – acetate. Green bars indicate higher DMS reactivity in methanol, whereas orange bars indicate higher DMS reactivity in acetate. Select bars are annotated for reference.

A different region of note in 23S rRNA is helix 101 near the 3′-end of the structure. This helix has several highly reactive nucleotides in double stranded regions (Supplemental Fig. S11, S12). Examination of *E. coli* 3D structure does not show any sign of accessibility of the equivalent nucleotides; however, this region is not covered by proteins, giving potential for breathing and higher reactivity.

Overall, reactivity for all three rRNAs in *M. acetivorans* was very similar in acetate and methanol growth media and consistent with our comparative analysis structures. However, a handful of specific, largely single-stranded nucleotides showed large differences in DMS reactivity between growth media.

## Conclusion

In this study, we developed a method for chemically probing the RNA from an organism from the *Archaea* domain of life, *M. acetivorans, in vivo*. This method was applied to cultures grown in different carbon and energy sources and resulted in high quality samples ready for NGS library preparation, specifically Structure-seq2 libraries. We also generated comparative structures of *M. acetivorans* 5S, 16S, and 23S rRNAs and then mapped the *in vivo* DMS reactivity values to them. These results indicated that the rRNA accessibility is consistent with the comparative structures and is very similar between carbon and energy sources. We did identify a few sites in each rRNA where DMS modification changed between acetate- and methanol-grown samples. These changes appear to be highly localized as the secondary structure of the surrounding motif does not change. The molecular basis for the changes in rRNA reactivity with growth media will require further study but could be due to changes in protein binding or rRNA modification (Kaberdina et al. 2009; Genuth and Barna 2018; Samir et al. 2018; Sun et al. 2021).

Our study opens the door to answering new and exciting questions surrounding RNA structure in the domain *Archaea*. Probing of rRNA expedites experiments on other RNAs including mRNA (Ding et al. 2014), tRNA (Yamagami et al. 2022), and novel ncRNAs, as well as other archaea organisms and chemical reagents (Bevilacqua and Assmann 2018; Mitchell et al. 2019a).

## Data Availability Statement

Sequencing data, all data for figures, supplementary figures, tables, and supplementary tables are available at GEO GSE229536.

## Acknowledgements

We thank the Genomics Core Facility at Pennsylvania State University for assistance with NGS experiments. We also thank Professors Sarah Assmann, Paul Babitzke, Timothy Meredith, and Scott Showalter for review and feedback on an early version of the manuscript. This work was supported by National Institutes of Health Grant R35-GM127064 (P.C.B), Pennsylvania State University Eberly College of Science Stanley R. Person Graduate Fellowship in Molecular Biology (A.M.W), and the Division of Chemical Sciences, Geosciences, and Biosciences, Office of Basic Energy Sciences of the U.S. Department of Energy through grant DE-FG02-95ER20198 MOD16 (J.G.F.).

## Footnotes

1 5S - https://crw2-comparative-rna-web.org/data/conservation_diagram/cons.5.a.Archaea.pdf; 16S - https://crw2-comparative-rna-web.org/data/conservation_diagram/cons.16.a.Archaea.pdf; 23S – 5’ https://crw2-comparative-rna-web.org/data/conservation_diagram/cons.23.a.Archaea.5.pdf, 23S – 3’ https://crw2-comparative-rna-web.org/data/conservation_diagram/cons.23.a.Archaea.3.pdf, single nucleotide frequency tables: 16S - https://crw2-comparative-rna-web.org/nucleotide-frequency/16s-rrna-model-single-base-frequency/; 23S - https://crw2-comparative-rna-web.org/nucleotide-frequency/23s-rrna-model-single-base-frequency/ base pair frequency tables: 5S - https://crw2-comparative-rna-web.org/nucleotide-frequency/5s-rrna-model-base-pair-frequency-tables; 16S - https://crw2-comparative-rna-web.org/nucleotide-frequency/16s-rrna-model-base-pair-frequency-tables/; 23S - https://crw2-comparative-rna-web.org/nucleotide-frequency/23s-rrna-model-base-pair-frequency-tables/

## Notes

### Competing Interest Statement

The authors have declared no competing interest.

